# A framework for the modular and combinatorial assembly of synthetic gene circuits

**DOI:** 10.1101/611574

**Authors:** Javier Santos-Moreno, Yolanda Schaerli

## Abstract

Synthetic gene circuits emerge from iterative design-build-test cycles. Most commonly, the time-limiting step is the circuit construction process. Here, we present a hierarchical cloning scheme based on the widespread Gibson assembly method and make the set of constructed plasmids freely available. Our two-step modular cloning scheme allows for simple, fast, efficient and accurate assembly of gene circuits and combinatorial circuit libraries in *Escherichia coli*. The first step involves Gibson assembly of transcriptional units from constituent parts into individual intermediate plasmids. In the second step, these plasmids are digested with specific sets of restriction enzymes. The resulting flanking regions have overlaps that drive a second Gibson assembly into a single plasmid to yield the final circuit. This approach substantially reduces time and sequencing costs associated with gene circuit construction and allows for modular and combinatorial assembly of circuits. We demonstrate the usefulness of our framework by assembling a double-inverter circuit and a combinatorial library of 3-node networks.

---

Synthetic biology relies on a design-build-test process. Decreasing costs of DNA synthesis have favored the construction of increasingly complex synthetic gene circuits^1-2^. However, this is often accompanied by a rise in time spent for molecular cloning, which can actually account for a substantial fraction of researchers’ activity. Due to technical limitations in the number of fragments that are efficiently assembled in a single reaction, complex gene networks usually cannot be constructed in a single step. Sequential cloning of circuit sub-components becomes tedious and time-consuming with increasing circuit complexity. Therefore, fast and efficient DNA assembly schemes are needed more than ever.

For most of current cloning methods a trade-off exists between modularity and seamless assembly. Golden Gate cloning^3^ and Gibson assembly^4^ are two popular cloning schemes that illustrate this trade-off. Golden Gate cloning and its derivatives, as well as other restriction-based methods such as the BioBrick standard^5^, offer modular assembly of parts and the possibility to easily build combinatorial libraries. However, the assembly process often leaves behind “scar” sequences between adjacent parts, which can affect the performance of the final construct, especially if present in coding sequences (CDSs) or untranslated regions (UTRs). Besides, these methods require a substantial *a priori* effort to obtain the required set of modular parts in the appropriate format. On the other hand, Gibson assembly and its derivatives, as well as other overlap-directed DNA assembly methods (e.g. SLIC^6^, SLiCE^7^, CPEC^8^), allow for seamless cloning. Nevertheless, this comes at the cost of modularity, since the interface between two assembled parts is usually unique and needs to be re-designed for each new combination. The need to design and synthesize unique overlapping sequences increases the assembly cost and delays the design-build-test cycle, and it makes these approaches less suitable for combinatorial assembly.

Recently, efforts have been made to address this trade-off, for example in the form of a modular variant of Gibson assembly known as MODAL^9^ or the Start-Stop assembly^10^ method that aims at modifying Golden Gate for seamless cloning. However, these new approaches are not exempt from the modularity-seamless trade-off, and thus increased modularity intrinsically requires linker sequences between parts^9^, and similarly a lack of between-part linkers comes at the cost of modularity^10^. These constraints might limit the widespread adoption of these assembly methods as compared to the parental Gibson and Golden Gate approaches.

For the assembly of complex gene circuits, a hierarchical cloning scheme is probably the best choice in terms of efficiency and speed. Cloning all parts in parallel within a single reaction soon becomes unrealistic as the number of parts increases. A sequential scheme divides the process in consecutive steps, which becomes lengthy as the number of steps increases. The strength of hierarchical cloning schemes relies on the division of the assembly process into discrete steps or levels, in which parallel assembly reactions are performed within a given level before their products are used as building blocks for the next level. For instance, MoClo^11^ is a hierarchical extension of Golden Gate in which parts such as promoters, UTRs, CDSs and terminators are first assembled into transcriptional units, which are subsequently combined into multigene constructs. 3G assembly^12^ combines Golden Gate and Gibson assembly in a hierarchical scheme that allows for single-day construction of complex circuits.

Despite modularity and hierarchical assembly of Golden Gate variants, Gibson assembly remains the most popular cloning method among synthetic biologists^13^. The reasons behind this may include a need or preference for seamless assembly, or the complex design and implementation of Golden Gate schemes, which can be daunting especially during the initial setup.

Here, we present a hierarchical cloning scheme based on the widespread Gibson assembly method. Starting from basic parts such as promoters, regulators, regulator binding sites or coding sequences, we first assemble transcriptional units (TUs) into intermediate plasmids. Next, we combine transcriptional units to clone gene circuits into the final recipient vector. Our hierarchical assembly scheme allows for modular and combinatorial assembly of gene circuits in only two steps in *Escherichia coli (E. coli)*. We demonstrate its usefulness by assembling a double-inverter circuit with almost 100% accuracy and a combinatorial library of 3-node networks.

## RESULTS AND DISCUSSION

First, we designed a multiple cloning site (MCS) for the construction of 3-node gene regulatory networks, and cloned it into a plasmid containing AraC, P_BAD_, a ColA replicon and kanamycin resistance^14^ to yield pCempty (Figure 1). The MCS contains three “slots” flanked by unique restriction enzyme (RE) sites for the insertion and removal of transcriptional units. Additionally, each slot contains an extra internal RE site, resulting in a total of three unique RE sites per slot. This allows us to keep a part permanently in the plasmid (e.g. a fluorescent reporter) and easily add extra parts up-or downstream. Slots are followed by strong Rho-independent transcriptional terminators to avoid read-through, and isolated from each other by biologically neutral 200bp spacers (designed with R2oDNA^15^) to minimize compositional context. The RE sites are separated by unique short spacers (12 or 30bp) that are lost during standard column purification of DNA, thus avoiding re-ligation. To allow for the hierarchical combinatorial assembly of networks with three transcriptional units, we split the MCS in three units, each harboring a slot, and each unit was cloned into plasmids with ampicillin resistance, giving the starting plasmids pTU-Aempty, pTU-Bempty and pTU-Cempty (Figure 1).

**Figure 1.**
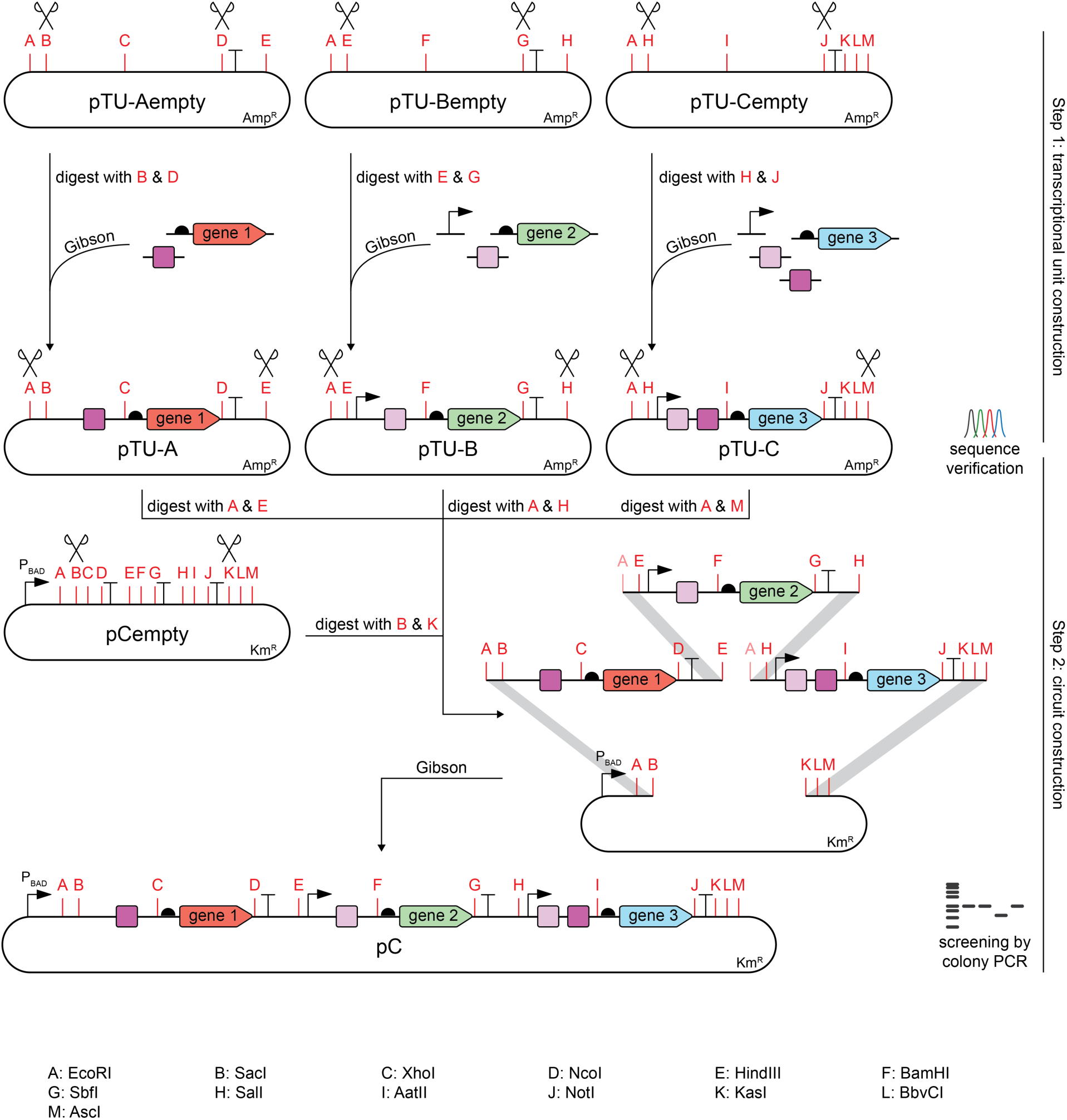
Schematic representation of a 3-node circuit constructing framework. The assembly occurs in two steps: during step 1 the individual transcriptional units (TUs) are assembled from their constituent parts into the intermediate plasmids (pTU-A, pTU-B and pTU-C), and in step 2 all the transcriptional units are combined into the receiver vector pCempty to yield the plasmid with the final circuit (pC). Both steps rely on RE digestion for vector linearization and on Gibson assembly for the directional, seamless cloning, and both allow for modular and combinatorial assembly. Note that during step 2 the EcoRI sites in TU-B and TU-C are not reconstituted in the final construct pC. Sanger sequencing verification is required at the end of step 1, but not after step 2. Bent arrows: promoters; squares: regulator binding sites; pointed rectangles: genes; semicircles: ribosome binding sites (RBS); and “T”-s: transcriptional terminators.

To construct a 3-node circuit, we carry out two steps (Figure 1). In step 1, the starting plasmids pTU-Aempty, pTU-Bempty and pTU-Cempty are opened with the appropriate REs, and the parts that will compose the transcriptional units are inserted using Gibson assembly. As a result, we obtain intermediate plasmids pTU-A, pTU-B and pTU-C, each one carrying one of the transcriptional units of the final circuit. Step 1 is modular in the sense that different promoters can be combined with different operators and coding sequences. This step also allows for the combinatorial cloning of a library of parts. Importantly, it is also possible to keep specific parts, such as fluorescent reporters, permanently in the starting plasmids, so that only variable parts (e.g. promoters and operators) need to be cloned between the remaining restriction sites.

In step 2, all three intermediate plasmids are digested with EcoRI plus an additional RE (HindIII for pTU-A, SalI for pTU-B and AscI for pTU-C), while the receiver vector pCempty is digested with SacI and KasI. As a result, transcriptional units A, B and C are flanked by overlapping regions that drive their directional A-B-C assembly into the linearized pCempty receiver plasmid. Digestions are column-purified and assembled in a Gibson reaction, resulting in the pC plasmid harboring the final user-designed circuit. Importantly, intermediate pTU-A, pTU-B and pTU-C vectors carry a different selection marker (Amp^R^) than the receiver pCempty plasmid (Km^R^) to avoid the recovery of non-digested/re-ligated pTU vectors at the end of step 2. Like step 1, step 2 is also modular (any pTU-A can be combined with any pTU-B and any pTU-C) and allows for the combinatorial cloning of transcriptional unit libraries. Each of the steps is performed in a single day; considering the time needed for assembly, bacterial growth and construct screening, a new synthetic gene circuit can be obtained from its basic constituent parts within 5 days. Once a library of TUs is available, composed circuits are assembled and screened within 24 hours.

To demonstrate the usefulness of our assembly method, we decided to assemble a 3-node gene regulatory network. The circuit is activated by arabinose (Ara) and deploys a double-inverter logic in which the first node (N1) represses the second node (N2), which subsequently represses the third node (N3) (Figure 2A and 2B). Repression is implemented through CRISPR interference (CRISPRi)^16^, for which a second plasmid carrying dCas9 (pJ1996_v2) is needed. This second plasmid also carries Csy4, a CRISPR endonuclease that recognizes and cleaves short RNA sequences^17^. We use csy4 recognition sites to “isolate” functional parts transcribed together within the same TU, so that once transcribed and physically separated by Csy4 they can act independently of each other. All three nodes carry a fluorescent reporter (mKO2, sfGFP and mKate2)^18-20^ tagged with a degradation tag (MarA, MarAn20 and RepA70^21^, respectively). N1 is under the control of a P_BAD_ promoter and produces a singleguide RNA (sgRNA-Z) that represses N2 by binding down-stream of the constitutive promoter (BBa_J23150). N2 in turn produces sgRNA-Y that represses expression of N3 controlled by the BBa_J23100 constitutive promoter. Thus, when subjected to increasing concentrations of Ara, N1 levels should rise while N2 levels decrease and N3 levels, in turn, increase.

**Figure 2.**
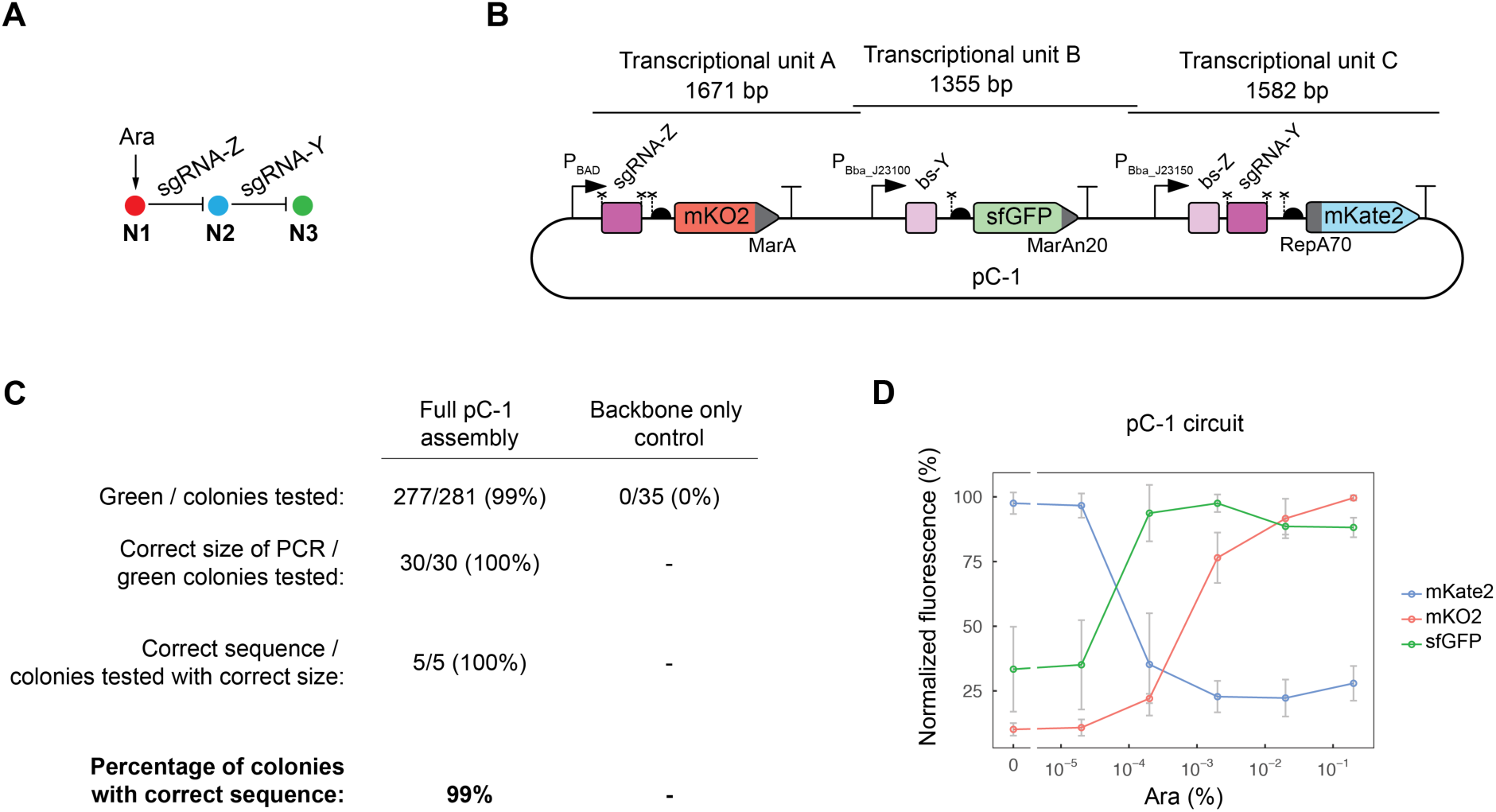
Implementation of the 3-node assembly strategy. A. The double-repression logic displayed by the 3-node circuit is depicted. B. Main relevant features of its implementation in pC-1, with the length of transcriptional units A, B and C shown. Bent arrows: promoters; squares: sgRNA binding sites; rectangles: sgRNAs; crosses: csy4 recognition sites; semicircles: RBSs; pointed rectangles: reporter genes with degradation tags; and “T”-s: transcriptional terminators. C. Accuracy of pC-1 assembly from its constituent single-node plasmids, as well as a control for backbone re-circularization. Three readouts with different throughput were used for determining accuracy: the fraction of green colonies (indicative of sfGFP presence), the fraction of green colonies carrying an insert of the correct length, and the fraction of colonies carrying a good-sized insert that showed no change at the nucleotide sequence level. Combining these data, we calculated the percentage of transformants that carried the correct sequence. Data based on two independent experiments. D. Assessment of the behavior of six non-sequenced transformants carrying the pC-1 circuit. Fluorescence levels in response to different arabinose concentrations indicate that all six transformants bear the pC-1 circuit that displays the expected double-repression logic. Mean and s.d. from six biological replicates.

In step 1, we used pTU-Aempty, pTU-Bempty and pTU-Cempty variants with fluorescent reporter genes (mKO2, sfGFP, mKate2 in pTU-A-005, pTU-B-005 and pTU-C-005, respectively) already present within the slots. We digested the three plasmids and inserted promoters, binding sites and sgRNAs upstream of the reporters using Gibson assembly. The sgRNAs were amplified from a storage vector and the promoters and binding sites were amplified from oligonucleotides. We used the MODAL strategy^9^ to facilitate the modular insertion of parts in step 1, but this is not a prerequisite. In step 2, the resulting intermediate plasmids containing the transcriptional units were digested with appropriate REs (Figure 1) and assembled into a RE-linearized receiver vector (the pCempty variant pC-0) to give pC-1 (Figure 2B).

Since only colonies incorporating TU-B should show green fluorescence, we used a blue light transilluminator for a pre-screen for green colonies. 99% of the colonies were green (Figure 2C), indicating that they contained TU-B. We performed colony PCR on green colonies using primers that anneal to the backbone and amplify all three nodes and observed that all (100%) green colonies carried the expected A-B-C insert (Figure 2C). Among those with the expected insert, five were randomly chosen for Sanger sequencing and all contained the correct sequence (100%). In summary, starting from a plate containing transformant colonies and in the absence of any pre-screening, the chances of picking a colony with the intended sequence were close to 100% (Figure 2C).

When building circuits of long sequence length, their verification through Sanger sequencing can become expensive and time-consuming. The cloning scheme presented here reduces sequencing-related time-and money-costs, since sequence verification is only needed after step 1 – at the intermediate plasmid level (Figure 1). Once an intermediate plasmid has been sequence-verified, it can be used repeatedly to build multiple circuits. In our experience, most of the mutations come from synthesized oligonucleotides used as ssDNA templates to generate short parts (e.g. promoters) or as primers, and only a small fraction results from high-fidelity PCR amplification. Gibson assembly uses a high-fidelity DNA polymerase to fill the short 3’ overhangs, and thus the probability of mutations arising during Gibson is low^4^. Since step 2 does not involve any PCR, the accuracy of the sequence of the final constructs is virtually 100%. Sequencing of the final construct is thus not necessary if the presence of all the TUs can be confirmed by other means (e.g. by colony PCR) (Figure 2C).

To confirm that this assembly framework can be used to efficiently build complex circuits that behave as intended with only one sequencing round (at the end of step 1), we assessed the behavior of colonies carrying the double-inverter pC-1 circuit. Briefly, plasmid DNA was extracted from six random pC-1 transformants verified by colony PCR, and transformed into *E. coli* MK01^22^ containing the plasmid carrying dCas9 and Csy4 (pJ1996_v2). Cells were grown in the presence of different Ara concentrations, and fluorescence was recorded. All six cloned exhibited the expected circuit behavior consisting of a rise in mKO2 levels with increasing Ara concentration, concomitant with an mKate2 decrease and a sfGFP increase (Figure 2D).

We next sought to investigate whether our method could be used to assemble combinatorial libraries of 3-node networks. The potential for combinatorial assembly is a desirable feature of DNA assembly methods, since it enables the building of large numbers of constructs without the need to design and build them individually. These libraries can then be rapidly tested provided that an appropriate high-throughput screening method is put in place^23-24^.

We set up a one-pot step 2 reaction in which we added 3 different pTU-A variants, together with 3 pTU-B and 3 pTU-C, which could result in a maximum of 27 different constructs (Figure 3A). Among these, some constructs may display varying repressive interactions between the three nodes, including the double-inverter (Figure 2), while others may lack any promoter and connectivity for some of the nodes. We determined the identity of 24 randomly picked transformants through Sanger sequencing, which revealed the presence of 9 different topologies, indicating the intended exploration of design space (Figure 3B). We measured fluorescence levels of the 9 different topologies and verified that they behave as expected according to their topology (Figure 3B and C). Therefore, our method provides a powerful platform for combinatorial assembly of gene circuits.

**Figure 3.**
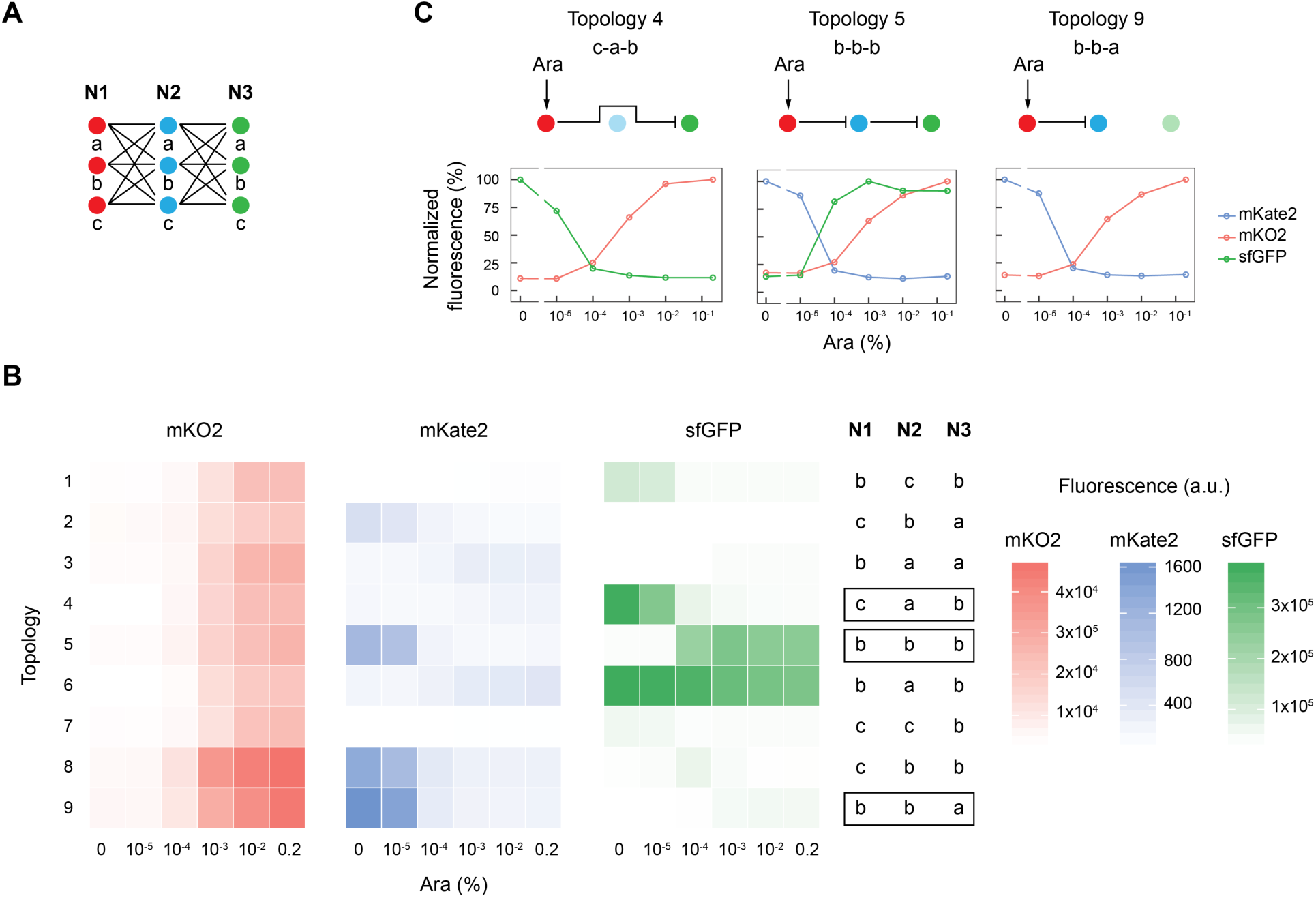
Combinatorial assembly of 3-node networks. A. Schema depicting all 27 different constructs that could result from the combinatorial assembly of three different N1 nodes, together with three N2 and three N3. B. Fluorescence levels of the 9 different topologies characterized in response to six arabinose concentrations. The identity of each construct is indicated on the right. C. Three topologies are shown in detail. The construct identity and the network connectivity are indicated, and normalized fluorescence levels are plotted below. N1 is always under the control of P_BAD_ promoter. N2 and N3 are either under the control of constitutive promoters (strong shading) or lack any promoter (weak shading).

We provide a set of 18 *E. coli* starting and receiver plasmids freely *via* Addgene upon manuscript publication (# 124409-124426; Table S1). In addition to the plasmids used for the examples described here, it also contains plasmids with alternative resistance markers, origins of replication and without AraC and P_BAD_ promoter. Our plasmids are designed for 3-node circuits, but the framework could easily be extended to host more nodes. Moreover, the framework is not restricted to gene circuits, but could also be used to assembly other synthetic systems, such as metabolic pathways.

In summary, we present here a time-and cost-efficient cloning scheme based on the popular Gibson assembly method that allows for modular and combinatorial assembly of synthetic circuits with an almost 100% accuracy. We believe that it is thus of interest for other synthetic biologists.

## METHODS

### DNA fragment preparation

sfGFP was amplified from pET-23100-lacO(SymR+1)-GFP-LVA plasmid^14^. mKO2 was purchased as a gBlock from IDT. mKate2 was amplified from plasmid pLPT107^25^, which was a gift from Johan Paulsson (Addgene plasmid # 85525). Csy4 was amplified from plasmid pCsy4^26^, which was a gift from Adam Arkin & Stanley Qi (Addgene plasmid # 44252). dCas9 was amplified from pJWV102-dCas9sp plasmid, which was a gift from Jan-Willem Veening^27^. Protein coding genes and sgRNAs^28^ were stored in a pJET1.2/blunt vector (Thermo Scientific) and amplified using KOD high-fidelity polymerase (Merck). For promoters and sgRNA binding sites, oligonucleotides were used as templates for KOD PCRs. All parts carried the same Prefix(CAGCCTGCGGTCCGG)and Suffix (TCGCTGGGACGCCCG) sequences for modular Gibson assembly using MODAL ^9^. Forward and reverse primers annealed to Prefix and Suffix sequences, respectively, and added unique linkers to the parts. Primers (desalted) were purchased from Microsynth or Sigma-Aldrich. All annealing steps were performed at 60°C. PCR amplifications were column-purified using the Monarch PCR & DNA Cleanup Kit (NEB) following manufacturer’s instructions. A list of primers can be found in the Supporting Information.

### DNA assembly and transformation

All digestions were performed for 1-2h at 37° with REs purchased from NEB, and purified using the Monarch PCR & DNA Cleanup Kit (NEB). Note that a complete digestion of receiver vectors in both step 1 and 2 is key to avoid the recovery of empty backbones. For step 1, plasmids were linearized using two REs (Figure 1) and PCR-amplified parts were assembled into the linear vectors using the NEBuilder HiFi DNA Assembly Master Mix (NEB) for 1h at 50°C, using a 2:1 insert:vector ratio or a 5:1 ratio when the insert was smaller than 250bp. For step 2, plasmids were digested as depicted in Figure 1, and purification was performed as mentioned above. Gibson assembly was performed with the NEBuilder HiFi DNA Assembly Master Mix (NEB) for 3h at 50°C using 50ng of receiver vector and a 2:1 insert:vector ratio. For the combinatorial assembly, variants of the same transcriptional unit were pooled for insert:vector ratio calculation. Of note, incubation of the Gibson reaction for shorter periods (1h) also results in very high assembly efficiency. 1µl of non-purifed Gibson reaction was transformed into 50µl of electrocompetent NEB5α cells, and 2/5 of them were plated onto selective agar plates.

### Colony PCR

Colony PCRs were run using Taq polymerase (NEB) with Thermopol buffer. All annealing steps were performed at 54°C. PCR products with the expected size were column-purified with the Monarch PCR & DNA Cleanup Kit (NEB) and Sanger sequenced by GATC-Eurofins or Microsynth. Transformants with the correct sequences were grown in 5ml selective lysogeny broth (LB), and plasmid DNA was extracted using the QIAprep Spin Miniprep Kit (QIAGEN).

### Fluorescence measurements

MK01^22^ electrocompetent cells were transformed with pJ1996_v2 plasmid (bearing dCas9 and csy4) as well as with pC-1 or a pC plasmid derived from the combinatorial assembly of 3-node networks (Figure 3). Single colonies were used to inoculate 2ml of selective LB, and after ∼6h cells were centrifuged at 4000rcf and resuspended in selective EZ medium (Teknova) with 0.4% glycerol as carbon source. 120µl of 0.05 OD600 bacterial suspensions were added per well on a 96-well CytoOne plate (Starlab), and 2.4µl of the indicated concentrations of L-arabinose (Sigma) were added. Plates were incubated at 37°C with double-orbital shaking in a Synergy H1 microplate reader (Biotek). Fluorescence was determined after 6h (for double-inverter) or 16h (for combinatorial library) with the following settings: mKO2: Ex. 542nm, Em. 571nm; sfGPF: Ex. 479nm, Em. 520nm; mKate2: Ex. 588nm, Em. 633nm. Fluorescence levels were corrected for fluorescence signal in a blank sample and for bacterial autofluorescence using a strain with no reporter genes and divided by OD600 to correct for differences in bacterial concentration.

## ASSOCIATED CONTENT

### Supporting Information

The following Supporting Information is available free of charge on the publisher website:

List of plasmids available through Addgene (Table S1).

List of primers for assembly, colony PCR and sequencing (Table S2).

## AUTHOR INFORMATION

### Author Contributions

JSM and YS designed research. JSM performed experiments, analyzed data and prepared the figures. JSM and YS wrote the manuscript. All authors have given approval to the final version of the manuscript.

## ACKNOWLEDGMENTS

We thank Marc García-Garcerà for help with R language, and Florence Gauye and Léo Moser for excellent technical assistance. We also thank all Schaerli lab members for useful discussions, and Lance E. Keller for critical reading of the manuscript.

## ABBREVIATIONS

Ara: Arabinose.
CDS: Coding Sequence.
MCS: Multiple Cloning Site.
MODAL: Modular Overlap-Directed Assembly with Linkers.
RBS: Ribosome Binding Site.
RE: Restriction Enzyme.
sgRNA: single guide RNA.
TU: Transcriptional Unit.
UTR: Untranslated Region.

