## Supporting Information for "A framework for the modular and combinatorial assembly of synthetic gene circuits"

| Name | Addgene ID | Resistance | Ori | Intended use | Comments |
|---|---|---|---|---|---|
| pTU-A-000 | 124409 | Ampicillin | pBR322 | pTU-Aempty, step1 | Empty slot A |
| pTU-B-000 | 124410 | Ampicillin | pBR322 | pTU-Bempty, step1 | Empty slot B |
| pTU-C-000 | 124411 | Ampicillin | pBR322 | pTU-Cempty, step1 | Empty slot C |
| pTU-A-005 | 124412 | Ampicillin | pBR322 | pTU-Aempty, step1 (contains reporter) | Slot A, contains mKO2-MarA |
| pTU-B-005 | 124413 | Ampicillin | pBR322 | pTU-Bempty, step1 (contains reporter ) | Slot B, contains sfGFP-MarAn20 |
| pTU-C-005 | 124414 | Ampicillin | pBR322 | pTU-Cempty, step1 (contains reporter) | Slot C, contains RepA70-mKate2 |
| pTU-A-013 | 124415 | Spectinomycin | CloDF13 | pTU-Aempty, step1 | Empty slot A |
| pTU-B-013 | 124416 | Spectinomycin | CloDF13 | pTU-Bempty, step1 | Empty slot B |
| pTU-C-013 | 124417 | Spectinomycin | CloDF13 | pTU-Cempty, step1 | Empty slot C |
| pTU-A-010 | 124418 | Spectinomycin | CloDF13 | pTU-Aempty, step1 (contains reporter) | Slot A, contains mKO2-MarA |
| pTU-B-010 | 124419 | Spectinomycin | CloDF13 | pTU-Bempty, step1 (contains reporter) | Slot B, contains sfGFP-MarAn20 |
| pTU-C-010 | 124420 | Spectinomycin | CloDF13 | pTU-Cempty, step1 (contains reporter) | Slot C, contains RepA70-mKate2 |
| pC-0 | 124421 | Kanamycin | ColA | pCempty receiver plasmid, step 2 | Contains full MCS |
| pC-0_v2 | 124422 | Ampicillin | ColA | pCempty receiver plasmid, step 2 | Contains full MCS |
| pC-0_v3 | 124423 | Ampicillin | pBR322 | pCempty receiver plasmid, step 2 | Contains full MCS |
| pC-0_v4 | 124424 | Ampicillin | pBR322 | pCempty receiver plasmid, step 2 | Contains full MCS, lacks AraC & P(BAD) |
| pC-0_v5 | 124425 | Ampicillin | pSC101 | pCempty receiver plasmid, step 2 | Contains full MCS |
| pC-0_v6 | 124426 | Ampicillin | pSC101 | pCempty receiver plasmid, step 2 | Contains full MCS, lacks AraC & P(BAD) |

**Table S1**. Plasmids available on Addgene.

| Name | Sense | Annealing site | Purpose* | Sequence(5'-->3')** |
|---|---|---|---|---|
| PR 79 | For | upstream MCS (in P(BAD) region) | colony-PCR of TU-A and of whole (3-node) circuit, sequencing of TU-A | ACGGCGTCACACTTTGC |
| PR 80 | Rev | spacer 1 | colony-PCR and sequencing of TU-A | AACTGGGTGGGACTCGGC |
| PR 81 | For | spacer 1 | colony-PCR and sequencing of TU-B | GCGAGAACTGCTGCCTGG |
| PR 87 | Rev | spacer 2.5 | colony-PCR and sequencing of TU-B | CACTGAAAAGTCTACGGAACTGCTGAG |
| PR 86 | For | spacer 2.5 | colony-PCR and sequencing of TU-C | CTCAGCAGTTCCGTAGACTTTTCAGTG |
| PR 89 | Rev | downstream MCS | colony-PCR of TU-C and of whole (3-node) circuit, sequencing of TU-C | GGGCCGTTGCTTCACAACG |
| PR 72 | Rev | 5' end of mKO2 | colony-PCR and sequencing of TU-A (when mKO2 reporter is present) | TCATCTCTGGTTTGATAACCGAAACCAT |
| PR 68 | Rev | 5' end of sfGFP | colony-PCR and sequencing of TU-B (when sfGFP reporter is present) | GTAAACAGTTCTTCGCCTTTACGCAT |
| PR 71 | Rev | 5' end of mKate2 | colony-PCR and sequencing of TU-C (when mKate2 reporter is present) | GCCATGTTATTTTCTTCTCCTTTACTAACCAT |
| PR 14 | For | Prefix | Adds Linker_14 overlap to insert TU-A during Step 1 | AGGATAGATTCTGGAAACTTTACCGTCCGAGCTC**CAGCCTGCGGTCCGG** |
| PR 145 | Rev | Suffix | Adds ECK120029600 Terminator overlap to insert TU-A during Step 1 | GCGGTCTTAAGTTTTTTGGCTGAACCATGG**CGGGCGTCCCAGCGA** |
| PR 15 | Rev | Suffix | Adds Linker_0 overlap to insert TU-A during Step 1 (when mKO2 reporter is present) | ACGGAGAAGCCCTATCAACTGTTTATTGCTCGAG**CGGGCGTCCCAGCGA** |
| PR 16 | For | Prefix | Adds Spacer 1 overlap to insert TU-B during Step 1 | CGAAATCCCTGAAACTGAGACTGTAGAAAATAAGCTT**CAGCCTGCGGTCCGG** |
| PR 153 | Rev | Suffix | Adds L3S2P21 Terminator overlap to insert TU-B during Step 1 | CTCTTTTCTGGAATTTGGTACCGAGCCTGCAGG**CGGGCGTCCCAGCGA** |
| PR 19 | Rev | Suffix | Adds Linker_10 overlap to insert TU-B during Step 1 (when sfGFP reporter is present) | ACAGTGCTCAATCGTGTAGAAATCTCTTGGATCC**CGGGCGTCCCAGCGA** |
| PR 20 | For | Prefix | Adds Spacer 2.5 overlap to insert TU-C during Step 1 | AGAAGTATTGGTAATCGTTGAAAACTCAGTCGAC**CAGCCTGCGGTCCGG** |
| PR 147 | Rev | Suffix | Adds ECK120033737 Terminator overlap to insert TU-C during Step 1 | CGGGCTTTTTTCTGTGTTTCCGCGGCCGC**CGGGCGTCCCAGCGA** |
| PR 23 | Rev | Suffix | Adds Linker_11 overlap to insert TU-C during Step 1 (when mKate2 reporter is present) | CTTCAGTTTCCAACGACGAGTGTAATAGGACGTC**CGGGCGTCCCAGCGA** |

*TU, transcriptional unit

** **Prefix** and **Suffix** sequences are indicated in **green** and **red**, respectively (for MODAL assembly, Casini et al. 2014). Users can replace these sequences with the 5' and 3' extremes of their TUs.

**Table S2.** Primers for assembly, colony PCR and sequencing.
